# Disruption of opioid and mesolimbic dopamine dynamics and behavior upon intranasal delivery of opioids in mice

**DOI:** 10.64898/2026.10.22.761063

**Authors:** Kellie J. Ervin, Maria F. Ramirez, Adrienn G. Varga, Daniel W. Wesson

## Abstract

The endogenous opioid system – consisting of mu, delta, and kappa opioid receptors, along with naturally occurring peptides, is central for regulating reward processing, mood, motivation, and social attachment. Endogenous opioids, as well as opioid drugs that are administered clinically or are misused/abused, are potent modulators of several neurotransmitter systems in the brain, including dopamine (DA). Since the opioid system is dysregulated in neuropsychiatric conditions, and opioids are prevalently misused/abused to achieve altered affective and cognitive states, the understanding of the mechanisms of action of opioids is highly significant. Unfortunately, there is no pre-clinical approach for understanding the actions of intranasally (IN) administered opioids – a common route of administration by both individuals with opioid use disorder and a route used clinically to achieve reversal of opioid overdose. Here we combined reliable and precise delivery of opioids into the nasal cavity of mice together with measures of opioid and DA dynamics in the nucleus accumbens (NAc) with fiber photometry, or, with measures of behaviors known impacted by opioids following other routes of administration, including respiration, to demonstrate the outcomes of IN opioids in mice. We uncovered a prompt and prolonged increase in mesolimbic DA by both IN morphine and fentanyl and the restoration of DA to baseline levels by subsequent IN opioid receptor antagonism with naloxone. Simultaneously, IN morphine and fentanyl each resulted in a short-lasting increase in NAc opioid dynamics. Next, in separate mice, we found that IN opioids elicit rapid canonical opioid-induced behaviors, including respiratory depression, which were rapidly reversed by IN naloxone. Overall, this work establishes a sensitive approach for evaluating the neural circuitry underlying opioid use disorder in a manner which recapitulates IN opioid use with ecological validity. Moreover, our findings point towards a likely important role for mesolimbic DA in mediating opioid overdose reversal.

## Introduction

Many drugs of abuse and neuropsychiatric disorders are associated with a dysregulation in brain dopamine (DA) and endogenous opioid systems^1–6^. These systems are central for reward processing^7,8^, emotion^9^, motivation^10^, and the addictive properties of many drugs of abuse^11^. Due to these important roles of DA and endogenous opioids, understanding of the mechanisms of action of opioids and its subsequent modulation of DA is highly significant.

Through their interaction with endogenous mu, delta, and kappa opioid receptors^12,13^, opioids promote analgesia^14^ and elicit overdose—which is commonly due to individual differences in tolerance^15^ as well as accidental taking of higher doses^16^. Fatal overdose symptoms are largely attributed to opioid agonism of specifically the mu opioid receptor^17^. In addition, the brain’s DAergic neurons contribute to opioid use disorder and euphoria through their rewarding effects^18,19^, most significantly via the opioid-driven disinhibition of DAergic neurons projecting from the ventral tegmental area to the ventral striatum, including DA release into the nucleus accumbens (NAc)^20–22^.

The use of intranasal (IN) administered opioids has increased in clinical practice, delivering rapid analgesia and overdose reversal with speed and efficacy^23^. Additionally, the snorting of opiates has surged both through the lacing of commonly nasally inhaled drugs and through purposeful taking^24^. The nasal mucosa contains both high vascularization and permeability and also has a large surface area due to the turbinate structure – both of these characteristics allow for efficient drug absorption^25^. The IN route also avoids hepatic and gastrointestinal first-pass metabolism^26^. The ultra potent synthetic opioid fentanyl is extremely lipophilic, causes low nasal irritability, and possesses a rapid onset of action upon binding to the opioid receptor, making it highly effective upon IN delivery, in manners which may explain both its addictive properties as well as its lethality^27,28^.

Existing *in vivo* models of opioid effects do not allow for assessing the neurobiological, behavioral, or respiratory effects of opioids when taken IN. This gap limits our understanding of the mechanisms of actions of opioids on the CNS, and specifically the possible subsequent dysregulation of opioid and DA dynamics. Routes of opioid administration other than IN—such as intramuscular (I.M.), intraperitoneal (I.P.), and subcutaneous (S.C.)—are known to elicit canonical opioid behaviors in rodents, including increased locomotion^29^, the Straub Tail repsonse^30,31^, respiratory depression^32–36^, and several other specific behavioral indicators of opioid exposure^37–39^. The occurrence, or possible absence, of these behavioral effects due to IN opioids are not known. Further, the latency at which these behaviors may occur upon the delivery of IN opioids and whether possible behavioral changes can be overcome by opioid receptor antagonism are also not known.

Here we administered IN morphine, fentanyl, and the opioid receptor antagonist naloxone to mice and quantified behavioral, respiratory, and neural responses upon delivery. To accomplish this, we leveraged a precise and reliable microfluidic device developed by our group, that when implanted on the nasal bone of mice, allows for rapid delivery of drugs and leads to systemic effects^40^. This approach allowed for the seamless measurement of the effects of IN opioid delivery and its reversal, without the need to restrain or inject the mice which fundamentally disrupts ongoing behavior and alters arousal. Our results uncovered the outcomes of IN opioids on neuromodulator and neuropeptide dynamics and the results of naloxone recovery by harnessing genetically-encoded AAV biosensors for dopamine (rDA3.8)^41^ and opioid dynamics (deltaLight)^42^. In a separate set of experiments, we established that IN opioids elicit rapid canonical opioid-induced behaviors and respiratory depression, the Straub tail response, and hyperlocomotion. We also verified that IN fentanyl induces overdose symptoms and that IN naloxone can rapidly reverse these effects and lead to subsequent canonical naloxone-evoked behaviors. Together, this work establishes a sensitive approach for evaluating the neural circuitry underlying opioid use disorder in a manner which recapitulates nasal opioid use with ecological validity and also points towards a likely important role for mesolimbic DA in mediating opioid overdose reversal.

## Methods

### Animals

Male and female C57BL/6J mice originating from breeder stock at the Jackson labs (strain #000664; RRID:IMSR_JAX:000664, Bar Harbor, ME), were bred and maintained at the University of Florida. Mice were housed in a temperature-controlled vivarium on a 12:12hr light-dark cycle with *ad libitum* access to food and water. All behavioral testing occurred during the light cycle. Mice were single housed following a single surgical procedure for chronic implants. Animals were housed in standard shoebox sized plastic cages on a ventilated rack. Corncob bedding and a Nestlet^TM^ (Ancaes, Bellmore, NY) were in each cage along with manzanita sticks (Bio-Serv, Flemington, NJ) for enrichment. All experimental procedures were conducted in accordance with the guidelines from the National Institute of Health and were approved by the University of Florida Institutional Animal Care and Use Committee.

We used a total of 21 4-7 months old mice for all experimentation. A total of 9 mice (5 males, 4 females) were used for *in vivo* fiber photometry experimentation, tested across three separate cohorts. Out of these mice, one was excluded for not demonstrating adequate photometric signal and one was excluded from averaged traces, but not statistics, due to a large detachment artifact during recording. An additional total of 12 mice (7 males, 5 females) were used for behavioral and respiratory measures and were tested across three separate cohorts. Of the 12, one failed a nasal patency test and did not contribute data to any measures. Another mouse had a complication with its nasal implant in the middle of experimentation and did not contribute data to the final two days of testing. Two mice (one in the naloxone recovery experiment and one in the HD fentanyl experiment) were excluded from reported data due to being statistical outliers (Microsoft Excel; IQR outlier testing).

### Surgical procedures

For all surgical procedures, under aseptic conditions, mice were anesthetized with 2-4% Isoflurane in oxygen (IsoFlo®, Patterson Veterinary, Houston, TX) and mounted on a stereotaxic frame while body temperature was maintained on a 38°C heating pad underneath their body. The analgesic Meloxicam (20mg/kg, Patterson Veterinary, Houston, TX) was subcutaneously (S.C.) administered followed by the local anesthetic lidocaine (lidocaine, 3mg/kg, S.C., Patterson Veterinary, Houston, TX) to the scalp before incision and exposure of the skull. Upon completion of surgeries, the mice were removed from the stereotaxic frame and placed in a clean home cage with a heating pad to aid recovery. Mice were given Meloxicam (20mg/kg, S.C., Patterson Veterinary, Houston, TX) for a 3-day post-operative period following the surgery.

### Indwelling nasal access port surgeries

To achieve microfluidic delivery into the nasal cavity, we harnessed the nasal access port (NAP) which was recently established by our group^40^. For NAP implantation, the nasal bone was leveled by adjusting the stereotaxic incisor bar. Next, a 1mm diameter hole was drilled through the right nasal bone, 1mm anterior to the frontal/nasal fissure and 0.5mm lateral. These coordinates were selected to position the NAP’s stainless-steel infusion tube immediately upon the vascularized turbinates of the nasal epithelium in adult C57BL/6J mice^40^. Following drilling, the infusion tube NAP was lowered through the craniotomy, and secured with dental cement (Teets cold cure, Diamond Springs, CA) to the proximal nasal bone and remainder of the skull. After the cement dried, a removable aluminum cap with a magnetic insert was placed over the implant to protect it while not in use.

### AAV injections and fiber implant surgeries

For *in vivo* monitoring of DA and opioid dynamics, we injected adeno-associated viruses to transduce the rDA3.8m (for DA) and deltaLight biosensors into the NAc^42,41^. For this, craniotomies were drilled to specifically target the medial shell region of the nucleus accumbens (+1.25mm AP/0.8mm ML/-4.0mm DV). A pulled glass micropipette containing 200nL of combined AAV9-hSyn-rDA3m (2.9×10^12^GC/mL) and AAV9-hSyn-deltalight (6.42×10^12^ GC/mL), both acquired from Neurotools (UNC Chapel Hill), was slowly lowered into the NAc and the virus delivered unilaterally using a Nanoject III (Drummond Scientific, Broomall, PA) at a rate of 2nL/s. Following 10-minutes, the pipette was slowly withdrawn from the brain. Subsequently, a 400μm outer diameter, 0.39NA optical fiber terminating in a 2.5mm outer diameter ferrule was lowered to the same coordinates as the injection, and cemented to the skull, for later fiber photometry-based monitoring of rDA3.8m and deltaLight levels. Next, in this same surgery, each mouse was implanted with a NAP, ipsilateral to the biosensors and fiber, as described above. Animals were allowed to recover for at least 3 weeks prior to any fiber photometry to allow sufficient viral transduction.

### Patency testing

Patency of NAPs was tested a day prior to any behavioral testing occurred. Mice were briefly anesthetized under 2-4% Isoflurane in oxygen and a 1mL syringe attached to an injector was used to infuse 0.1mL of saline through the NAP to ensure a small amount of fluid could travel through the NAP and into the nasal cavity as evidenced by a visible fluid drop exiting the ipsilateral nostril.

### Drugs

Fentanyl citrate, morphine sulfate, and naloxone hydrochloride were each purchased in solid salt state from Millipore Sigma and diluted to desired concentrations using sterile saline. Concentration was calculated w/v and adjusted to achieve desired concentration per infusion based on body weight of the animal (mg/kg/inf). Due to the nature of delivering solutions into the nasal cavity of mice, the volumes must remain small to avoid asphyxiation and to minimize the possibility drug would enter the esophagus or pass out through the nostrils^40^. With this volume limit in mind, morphine concentrations could not exceed 75mg/mL due to the lesser solubility and potency of morphine compared to fentanyl and naloxone. Bodyweight-based dosing of drugs when applicable and also a timeline of the drug-delivery schedule within groups of mice are included in **Table 1** and **Table 2**, respectively.

**Table 1.** Drug concentration groups based on body weight for behavioral and respiratory experiments.

| *Weight Classes* |  | 18-23g | 28-33g | 33.1-38g |
| --- | --- | --- | --- | --- |
| IN morphine | Dose | 50mg/mL | 75mg/mL | 75mg/mL<sup>1</sup> |
|  | Number of animals | 4 | 6 | 1 |
| IN low dose fentanyl | Dose | 0.2mg/mL | 0.31mg/mL | 0.35mg/mL |
|  | Number of animals | 4 | 6 | 1 |
| IN high dose fentanyl | Dose | 1.5mg/mL | 2.3mg/mL | 2.6mg/mL |
|  | Number of animals | 3 | 3 | 1 |
| I.P. high dose naloxone | Dose | 0.2mL of<br>0.15mg/mL | 0.31mL of<br>0.15mg/mL | 0.35mL of<br>0.15mg/mL |
|  | Number of animals | 2 | 2 | 1 |
| I.P. naloxone | Dose | 30mg/mL | 46mg/mL | 53mg/mL |
|  | Number of animals | 4 | 5 | 1 |
| IN naloxone | Dose | 0.1mL of<br>6mg/mL | 0.15mL of<br>6mg/mL | N/A<sup>2</sup> |
|  | Number of animals | 2 | 2 | 0 |

*Note.* All drugs administered to each animal throughout the course of experimentation is in line with these groupings. Groups were formed based on $\pm 10\%$ body weight.
<sup>1</sup>Morphine sulfate dissolved in sterile saline has a max concentration of 75mg/mL due to solubility, so mice heavier than 33g had to be grouped with the 28.1-33g category despite precise body weight for morphine administration only.
<sup>2</sup>I.P. naloxone for the 33.1-38g weight class was not needed.

**Table 2.**
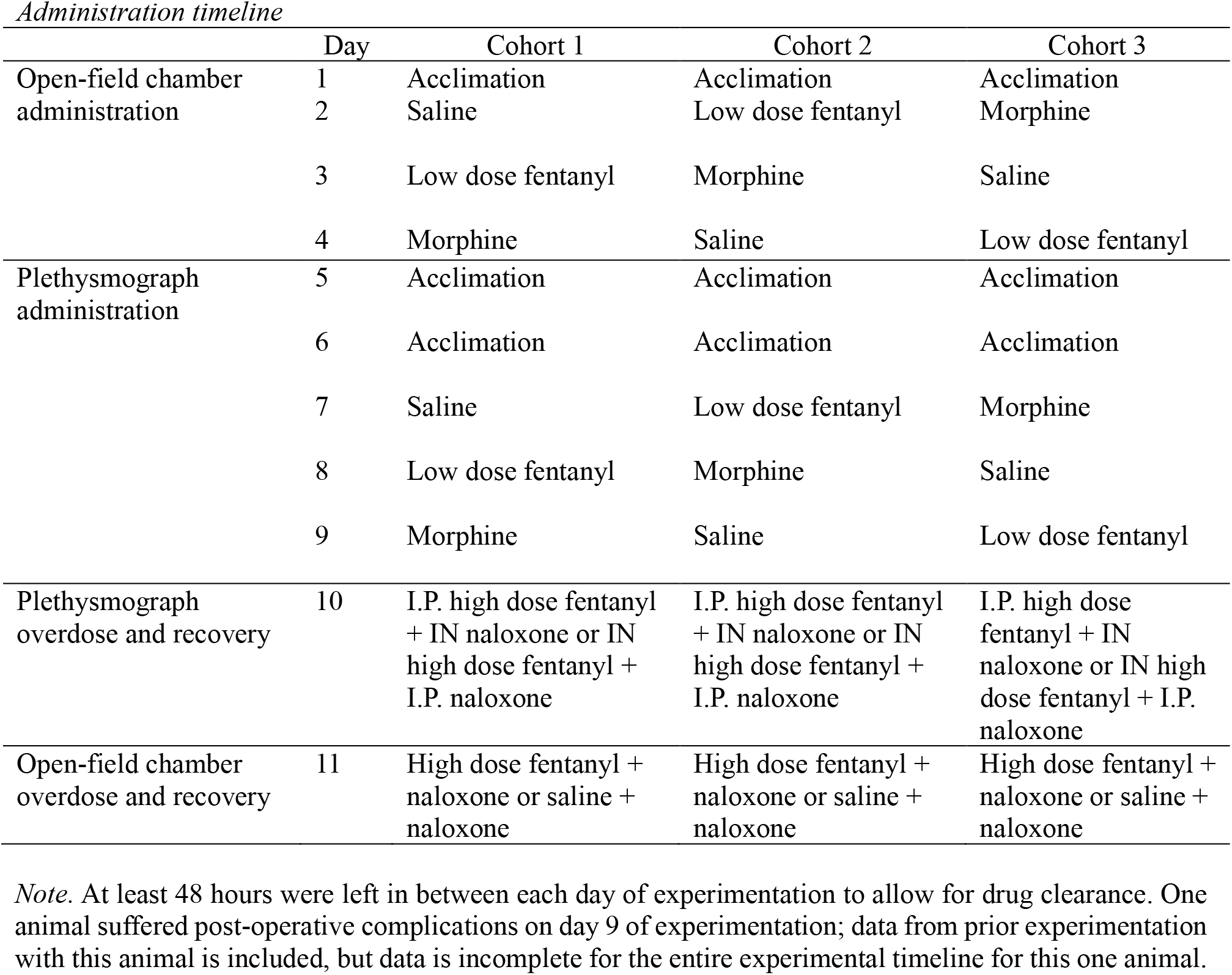
Drug administration timeline for three separate cohorts of mice during behavioral and respiratory experimentation.

### In vivo fiber photometry with IN drug delivery experimentation

560nm (rDA excitation wavelength, driven at 250 Hz), 465nm (DOR excitation wavelength, driven at 210 Hz), and 405nm (UV excitation wavelength, driven at 330Hz; control) light emitting diodes (Tucker Davis Technologies, Alachua, FL, RZ10x) were coupled to a fluorescent minicube (Doric lenses, FMC6) using 200μm, 0.22NA FCM optic fiber patch cords. Excitation and emission light were directed through a single 400μm, 0.57NA patch cord connected to the mouse by the implanted 2.5mm metal ferrule. Emission light was directed through the minicube and coupled to femtowatt photoreceivers to monitor GFP, RFP, and UV signals (Tucker Davis Technologies, Alachua, FL, RZ10x). The emission signals were low-pass filtered (2nd order), pulse demodulated, and digitized (1kHz) using a Tucker Davis Technologies processor. All parameters for acquisition of the photometry data were consistent across all groups. UV isosbestic channel was manually analyzed for major artifacts and subtracted from RFP and GFP channels to eliminate artifacts.

### Locomotor activity and behavioral testing

Locomotion and other behaviors were measured in a custom-made, open top, acrylonitrile butadiene styrene 30.48 × 30.48 × 30.48cm dark acrylic chamber (length × width × height). The mice were handled and habituated to the behavior room prior to experimentation. On Day 1, the animals were gently scruffed, the protective aluminum cap on the NAP was removed, and the mice were tethered to a flexible infusion line. The line was empty to allow habituation to the apparatus. On days 2, 3, and 4, sterile saline, a low dose of fentanyl (0.2mg/kg), and morphine (50mg/kg) were administered in a randomized fashion. The lines were backfilled with the treatment drug and a fluid-tight swivel connected to a 1mL syringe in a pump powered and controlled by an Arduino Uno microcontroller. The Arduino generated a pulse to trigger the pump to deliver 2µL infusions via the NAP into the nasal epithelium every 15s, for a total 20µL bolus. The mice were placed in the chamber for 5 minutes to acclimate each day tethered to the swivel and at the 5 min mark, they received infusions and locomotor and behavioral activity was recorded for 20 minutes post-infusion. At the end of the 20 minutes, mice were gently restrained, the tether was removed, and they were placed back in their home cage.

On day 11, some mice were tethered to a line backfilled with a high dose of fentanyl to begin measurement and some were tethered to a line backfilled with saline. The mice were placed in the chamber for 5 minutes to acclimate tethered to the swivel and at the 5 min mark, they received infusions and locomotor and behavioral activity was recorded for 15 minutes post-infusion. The line was then switched for one backfilled with naloxone, the mice received infusions, and behavior and locomotor activity was recorded for 10 minutes before the mice were removed from the apparatus and returned to their home cage.

All behavioral data were recorded using two video cameras (17.2 frames per second) located above and horizontal to the chamber. Locomotion was extracted from the videos off-line in ezTrack^43^. For all behavioral experiments, mice were allowed at least 48 hours of recovery in between testing days to allow for drug clearance.

### Assessment of breathing

To monitor breathing, mice were placed in whole-body plethysmograph (Data Sciences International, St. Paul, MN). On days 5 and 6, the animals were gently scruffed, the protective aluminum cap was removed, and the mice were tethered in the plethysmograph to allow for acclimation. The fluid delivery line was empty to allow habituation to the apparatus. On days 7, 8, and 9, sterile saline, a low dose of fentanyl (0.2mg/kg), and morphine (50mg/kg) were administered in a pseudorandomized fashion while the mice were inside of the plethysmograph. The mice were placed in the plethysmograph for 5 minutes to acclimate each day tethered to the swivel and at the 5 min mark, they received IN fluid infusions and behavioral activity was recorded for 20 minutes post-infusion. At the end of the 20 minutes, mice were gently restrained, the tether was removed, and they were placed back in their home cage.

On day 10, some of the mice were given an I.P. injection of a high dose of fentanyl (1.5mg/kg) and tethered to a infusion line that was backfilled with naloxone (30mg/kg) before being placed in the plethysmograph. A baseline measurement of 3 minutes was completed before naloxone was administered, and at least 13 minutes of respiration was recorded before the mice were removed from the apparatus and returned to their home cage. The other mice were tethered to a line that was backfilled with a high dose of fentanyl (1.5mg/kg) before being placed in the plethysmograph. Respiration was measured for at least 13 minutes before the mice were removed from the apparatus and given an I.P. injection of naloxone (30mg/kg) before being returned to their home cage. All lines were backfilled with the treatment and the swivel connected to a 1mL syringe in a pump powered and controlled as described in the previously mentioned experiments.

Respiratory transients were detected using a pressure transducer (Data Sciences International) and digitized at 300 Hz (0.1–12 Hz band pass; Tucker Davis Technologies, Alachua, FL), following 500x gain amplification (Cyngus Technology Inc, Southport, NC). Positive pressure of clean room air was continuously applied to the chamber using a stable-output air pump (Tetra Whisper, Blacksburg, VA). Inhalation peaks were detected offline in Spike2 (Cambridge Electronic Design, Cambridge, UK) and breathing frequency calculated by comparing the inter-breath intervals. For all respiratory experiments, mice were allowed at least 48 hours of recovery in between testing days to allow for drug clearance.

### Statistical Analysis

All data were processed using semi-automated steps whenever possible, including processing data from mixed treatment groups at the same time. For photometry and respiratory analyses, repeated measures ANOVAs were used. Behavioral scoring was not blind due to the nature of the experimenter also being the person who loaded drugs into the syringe pump, and all videos were scored by a single observer (K.J.E.) with inter-observer reliability spot-checking by D.W.W. Statistical comparisons of behavior were analyzed using one-way ANOVAs. All data are reported and plotted as mean ± SEM unless noted otherwise. Data from some individual mice that did not pass an outlier test (Microsoft Excel; IQR outlier test) are not included in figures nor in the statistical reporting and these instances are mentioned in the respective figure legends.

## Results

### IN opioids result in real-time disruption of endogenous opioid and dopamine dynamics in the brain

As a starting point, we sought to evaluate the impact of IN opioids and naloxone reversal on neural dynamics. For this, we used *in vivo* fiber photometry paired with genetically-encoded AAV biosensors for dopamine (DA, rDA3.8m)^41^ and the delta endogenous opioid receptor (DOR, deltaLight)^42^ in the reward system. We chose the deltaLight sensor specifically given its high affinity and relatively rapid temporal dynamics upon binding opioids^42^. Mice were injected with both sensors into the NAc and, several weeks later, implanted with a multimode optical fiber into the ipsilateral NAc as well as a nasal access port (NAP)^40^ for IN drug delivery. (**Fig. 1A**). We targeted the medial NAc shell specifically due to its innervation by the medial VTA (where opioid peptides bind and disinhibit DA signaling) and its role in the reinforcing aspects of opioids^44^. At least five days later to allow recovery, mice were tethered to for an optical patch cable photometry data collection and tethered to their NAP for IN drug delivery (including morphine, a high dose (HD) of fentanyl, naloxone, and a saline control; see Methods). On each recording session, each mouse was allowed to acclimate to the tethering and the behavioral chamber before drug delivery (**Fig. 1B**). All data presented herein originated from mice with confirmed histological placement of the AAV-based biosensors and the optical fiber (**Fig. 1C**).

**Fig 1.**
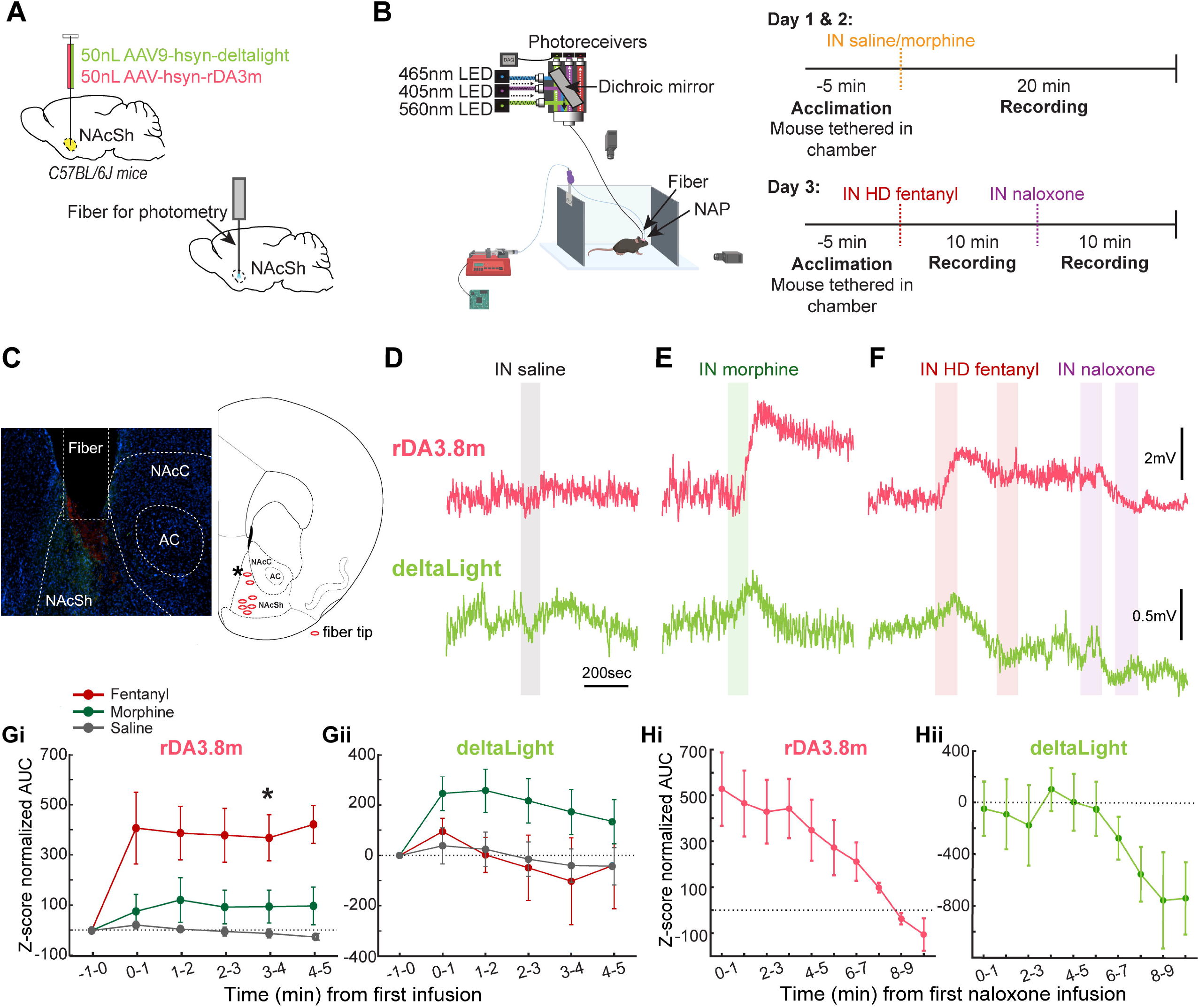
IN opioids result in real-time disruption of endogenous opioid and dopamine dynamics in the brain. **A.** Schematic of AAV injection and implantation of an optical fiber for fiber photometric collection of signals from the NAc of mice. **B.** Behavioral setup for simultaneous fiber photometry recording and drug delivery with experimental timeline. **C.** Anterior-posterior position of AAV and fiber placement, example fluorescent histology image of the NAc (green = deltaLight, red = rDA3.8, and blue = DAPI), and representative image of fiber placements for all animals included in study (coronal diagram, apx 1.7mm ant bregma). * represents the fiber placement of the example fluorescent image. rDA3.8 and deltaLight fiber photometry traces when IN saline (**D**), IN morphine (75mg/mL) (**E**), and IN HD fentanyl (2mg/mL) and IN naloxone (60mg/mL) (**F**) were delivered to the same animal. Averaged area under the curve (AUC) of all traces during administration of IN morphine (n = 7), saline (n = 7), and HD fentanyl (n = 4) in the red (**G**) and green (**H**) channels (baseline = 0). *=*p*<0.05, fentanyl vs saline at 3-4mins. Dashed horizontal line = normalized baseline. **I.** Averaged AUC of all traces during administration of IN naloxone after fentanyl-induced overdose of red and green channels (n = 4). Created in BioRender. Wesson, D. (2027) https://BioRender.com/7xd28hz.

As reflected by example traces from one mouse, IN delivery of both morphine and fentanyl resulted in a real-time increase in rDA3.8m levels (**Figs 1 E&F**). This elevation in rDA3.8m was considerably rapid (occurring prior to the end of the infusion period, <1min) and prolonged (lasting for several mins). Simultaneously, morphine and fentanyl also resulted in a similar rapid increase in deltaLight (**Figs 1E&F**). Across the population of mice, increases in rDA3.8m were apparent for both fentanyl and, to a lesser extent, morphine (rDA3.8m: condition (*F*(1.20, 4.80) = 7.70, *p* = 0.0383), time (*F* (1.66, 6.63) = 10.38, *p* = 0.0105), condition by time interaction (*F* (2.19, 8.75) = 4.50, *p* = 0.0430); deltaLight: condition (*p* = 0.9095), time (*p* = 0.2425), condition by time interaction (*p* = 0.8522); **Fig 1G & H**). Post hoc paired comparisons showed that the AUC was significantly higher under IN HD fentanyl administration than under saline as soon as the fourth minute after administration (fourth minute: *p* = 0.0416; fifth minute: *p* = 0.0178; **Fig. 1G**). The population-level result also indicates an increase in deltaLight by morphine, but not by fentanyl (**Fig 1H**).

In the same mice, within sessions where they received IN fentanyl, we subsequently delivered IN naloxone upon visual signs of overdose (*i.e.,* gasping, loss of limb mobility/sternal recumbency) to examine rDA3.8m and deltaLight dynamics during antagonism of opioid receptors. As shown by the example traces from a single mouse in **Figure 1F**, IN naloxone restored levels of rDA3.8m, which had risen upon IN fentanyl, back down to baseline levels (that being the level pre-fentanyl). Conversely, IN naloxone resulted in a further suppression of deltaLight levels to those even lower than following IN fentanyl. This reduction of rDA3.8m back to baseline and, separately, the further suppression of deltaLight signals were similarly observed across the population of mice, with significant differences in the rDA3.8m signal detected around 2 minutes post infusion and the deltaLight signal detected around 5 minutes post infusion (Repeated measures ANOVA; time: *F*(9,36) = 6.38, *p* < 0.0001; signal type [rDA3.8m or deltaLight]: *F*(1,4): 18.08, *p* = 0.013; rDA3.8m 0 min vs 2 min: *p* = 0.023; deltaLight 0 min vs 5 min: *p* = 0.016; **Figs 1I & J**). Together, these results indicate real-time disruption of endogenous DA and opioid dynamics in the brain upon IN opioids, and also demonstrate that IN naloxone can quickly reverse both DA and opioid disturbances post-fentanyl overdose.

### IN opioids elicit canonical opioid behaviors in freely-moving mice, including the Straub tail response

Next, in separate mice, we conducted a within-subjects study to assess the behavioral outcomes of IN opioids and the efficacy of IN naloxone to overcome opioid overdose. We monitored behavior in an open field chamber for video-based quantification of locomotor^29^, Straub tail^30,31^, and other behaviors that have been reported following opioid administration^37–39^ (**Fig. 2**). For this, all mice were implanted with a NAP to allow for IN drug administration and allowed to recover for at least one week. Afterwards, mice were tethered to a drug infusion line to their NAPs and placed in the open field chamber (**Fig. 3A**). Over subsequent days, the same mice were connected to the drug infusion tether and placed into a whole-body plethysmograph chamber for quantification of possible respiratory depression and its reversal by naloxone. As opposed to the across-mouse dosing in the photometry experiments above, the mice in these respiratory experiments received individualized dosing based on their weight to allow for assessing behavior and respiration, and their changes with specific dosages (**Table 1**). Mice were delivered the different drug treatments in a pseudorandom order to prevent a testing-order effect: with mice receiving saline, morphine, and low dose (LD) fentanyl on different days and in a Latin square type design (**Table 2**). At least 48 hours elapsed between testing days to allow a drug wash-out period. As a final testing condition, mice received near-lethal high-dose (HD) fentanyl, and we examined recovery of behavior following naloxone, on separate days (**Fig. 2C & D**).

**Figure 2.**
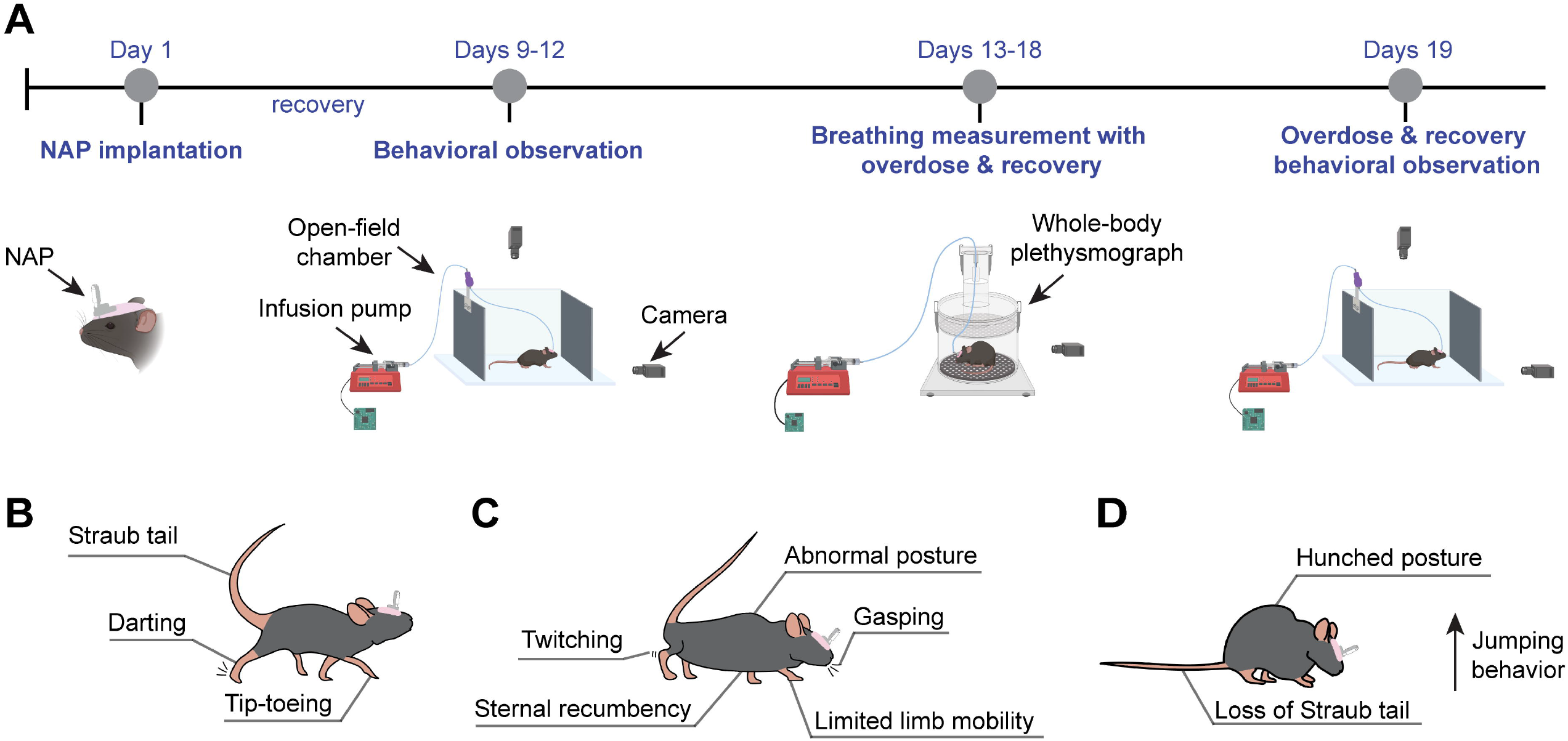
Experimental timeline and key behavioral measures. **A** Timeline of experimentation, including surgical implantation of a nasal access port (NAP) to allow IN drug delivery, behavioral and locomotive testing in the open-field chamber, and respiratory recordings in a whole-body plethysmograph. Within each testing condition (behavioral observation, breathing measurement with overdose and recovery, or behavioral observation with overdose and recovery) drugs were delivered in a pseudorandom order within each mouse (see **Table 2**). Each administration day allowed 5 minutes for the mouse to be acclimated to the tether and, following drug delivery, we acquired data for 20 minutes. Mice were given at least 48 hours for recovery in between each experimental day to allow for drug clearance before beginning the next experiment. Illustration of behaviors monitored during behavioral observation and measurement when IN saline control, IN LD fentanyl, and IN morphine were administered (**B**), during overdose measurement after administration of IN HD fentanyl (**C**), and during recovery measurement after administration of IN naloxone (**D)**. Created in BioRender. Wesson, D. (2027) https://BioRender.com/7xd28hz.

**Figure 3.**
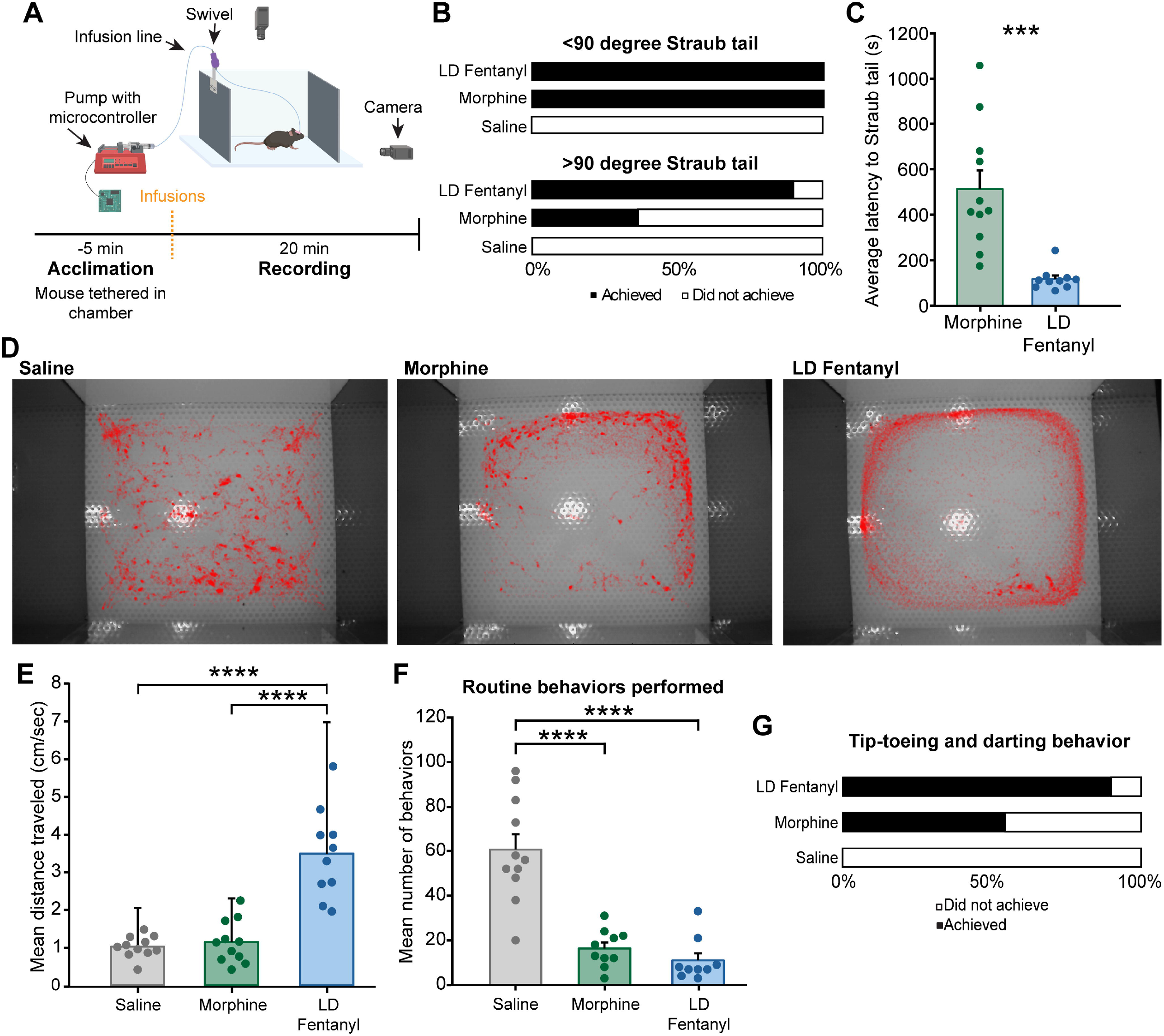
IN opioids elicit canonical opioid-related behaviors in freely-moving mice. **A.** Experimental setup of freely-moving chamber for behavioral measurement with IN delivery and experimental timeline. Drugs were automatically infused into the nose by means of a syringe pump controlled by a microcontroller while locomotion and behaviors were recorded by overhead and horizontal cameras. **B.** Proportion of mice achieving a Straub tail less than 90 degrees above the horizontal plane after various treatment administrations and proportion of mice achieving a Straub tail that was more than 90 degrees above the horizontal after various treatment administrations. **C.** The recorded latencies to achieving the Straub tail response after drug administration. ***=*p*<0.001. **D.** Motion tracing showing the locomotive patterns of the same mouse after receiving IN treatment in the open field chamber. **E.** The distance (cm/s) traveled by the mice depending on the treatment administration. **F.** The number of behaviors mice performed during the entire experimental session, including facial grooming, rearing, and itching. ****=*p*<0.0001. **G.** The proportion of mice that performed tip-toeing and darting behaviors (canonical opioid locomotive behaviors) during various treatments. Created in BioRender. Wesson, D. (2027) https://BioRender.com/7xd28hz.

We found that all mice given IN morphine and LD IN fentanyl displayed a mild Straub tail response, demonstrated by a lifting of the tail less than 90 degrees above the horizontal plane (**Fig. 3B**). IN LD fentanyl led to 90% of the mice displaying the extreme Straub tail, demonstrated by a lifting of over 90 degrees above the horizontal of the tail, yet morphine only resulted in an extreme Straub tail reaction in 40% of the mice (**Fig. 3B**). No Straub tail responses were observed following IN saline (**Fig. 3B**).

One advantage of delivering the opioids IN in this paradigm is that the drugs can be administered without any manual intervention of the experimenter (*viz*., scruffing). Therefore, we took advantage of this data set to also quantify the latency from IN infusion to changes in behavior **(Fig. 3C**). We found that IN morphine administration resulted in a relatively gradual latency to Straub tail response (513.54 ± 82.41 SEM). In contrast, IN fentanyl resulted in a prompt Straub tail response, even compared to morphine (117.80 ± 15.44; One-Way ANOVA, morphine vs LD fentanyl: *F* (1,19) = 20.28, *p* = <0.001; **Fig. 3C)**.

We further found that LD IN fentanyl resulted in hyperlocomotion. Both LD fentanyl and morphine elicited a locomotor pattern characterized by mice spending most of their time locomoting along the edges of the chamber (**Fig. 3D**). Mice receiving LD fentanyl traveled a greater distance than both the morphine (One-Way ANOVA, LD fentanyl vs morphine: *F* (1,19) = 33.81, *p* = <0.0001) and saline infused groups (One-Way ANOVA, LD fentanyl vs saline: *F* (1,19) = 44.08, *p* = <0.0001; morphine vs saline: *F* (1,20) = 0.41, *p* = 0.53l; **Fig 3E**).

Other behaviors were also impacted following IN opioids. We observed a significant decline in routine exploratory behaviors, including grooming and rearing, following morphine and LD fentanyl (One-Way ANOVA, morphine vs saline: *F* (1,19) = 32.51, *p* = <0.0001; LD fentanyl vs saline: *F* (1,19) = 35.85, *p* = 0.53l; **Fig. 3F**). Abnormal gait and paw placement behaviors were also observed, including darting and tip-toeing behaviors, with 83.3% of mice that received IN HD fentanyl displaying one or both behaviors, 90% of mice receiving IN LD fentanyl displaying one or both behaviors, and 54.5% of mice receiving IN morphine displaying one or both behaviors. None of these behaviors were observed following IN saline (**Fig. 3G**).

### IN opioids result in respiratory depression

On later testing days, the same mice used above were tethered to an infusion line for IN drug delivery and placed into a whole-body plethysmograph for measures of respiration (**Fig. 4A**). As expected by the extensive literature on respiratory depression in mice following peripheral administration (e.g., intra-peritoneal^32,36^), we found that IN morphine and LD fentanyl elicited mild respiratory depression (**Fig. 4B-D**). When the same mouse was compared between IN saline, morphine, and LD fentanyl administration days, there was a visible depression in respiratory frequency after morphine and fentanyl administration that was not seen with saline (**Fig. 4B-D**).

**Figure 4.**
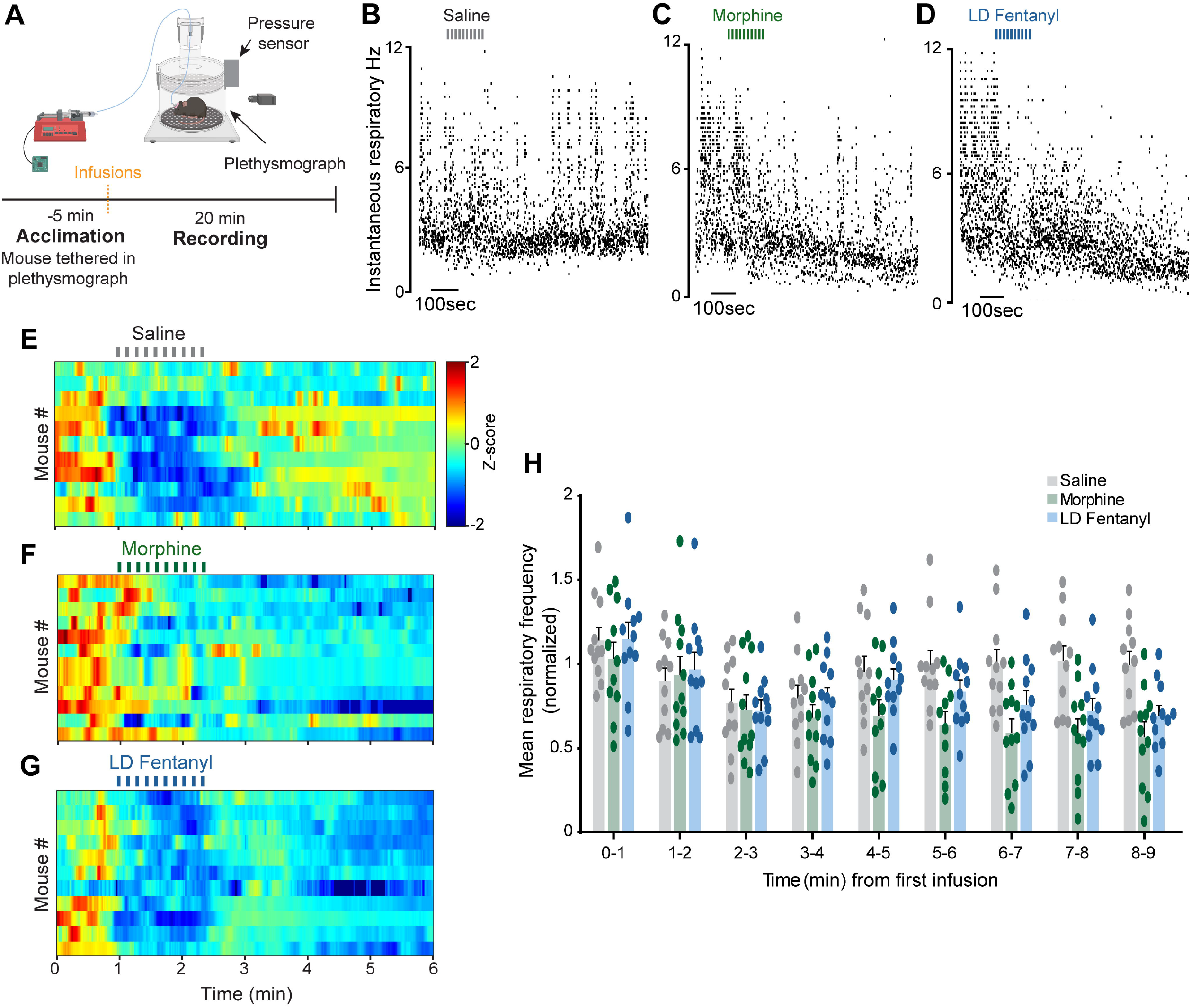
IN opioids result in respiratory depression. **A.** Whole-body plethysmography setup and experimental timeline. Drugs were automatically infused into the nose via a syringe pump controlled by a microcontroller while video was recorded by a side-view camera. **B.** Respiration recorded from a mouse in the plethysmograph when IN saline was administered. **C.** Respiration recorded from the same mouse in the plethysmograph when IN morphine was administered. **D.** Respiration recorded from the same mouse in the plethysmograph when IN low dose fentanyl was administered. Instantaneous frequency of respiration of all animals when IN saline (**E**), morphine (**F**), and low dose fentanyl (**G**) were administered. **H.** Instantaneous frequency of respiration demonstrated in 60-second bins for saline, morphine, and low dose fentanyl administration. Created in BioRender. Wesson, D. (2027) https://BioRender.com/7xd28hz.

All mice displayed respiratory depression after administration of IN morphine and LD fentanyl (**Fig. 4E-G**). IN morphine and LD fentanyl elicited a significant respiratory depression compared to the IN saline control (Repeated measures ANOVA at the interception of condition and time; *F*(16,160) = 7.61, *p* < 0.0001; **Fig. 4H**).

### Overdose symptoms after administration of high dose IN fentanyl

We next wanted to examine respiratory consequences upon IN HD fentanyl and the ability for IN naloxone to recover mice from an overdose of fentanyl. Because only one drug can be infused at a time through the nasal port, the mice were divided into two groups to address each of these goals, which notably were saved until the end of experimentation due to the risk of lethality from HD fentanyl. One subset of mice received IN HD fentanyl while in a plethysmograph to assess the inset of respiratory depression, to be later reversed by an I.P. injection of naloxone. A second subset of mice were overdosed with an I.P. injection of HD fentanyl prior to examining the ability for IN naloxone to overcome respiratory depression. Both subsets of mice were, on a subsequent day, monitored in an open field chamber for the ability of IN naloxone to overcome the behavioral effects HD fentanyl administered IN.

Upon IN HD fentanyl (**Fig 5A**), we observed extreme respiratory depression, which occurred about 70 seconds after the first infusion (**Fig. 5B**). During IN HD fentanyl-induced respiratory depression, little to no sniffing or otherwise alert breathing dynamics were observed (**Fig. 5C**). There was a severe decline in respiratory frequency over time, with respiration being significantly depressed as soon as 2 minutes post-infusion onset (Repeated measures ANOVA, *F*(8,24) = 46.371, *p* < 0.0001; 0 min vs 2 min: *p* = 0.0032; **Fig. 5D**). Most mice during this session experienced a gradual, yet dramatic, decrease in respiratory frequency upon IN HD fentanyl administration (**Fig. 5E**).

**Figure 5.**
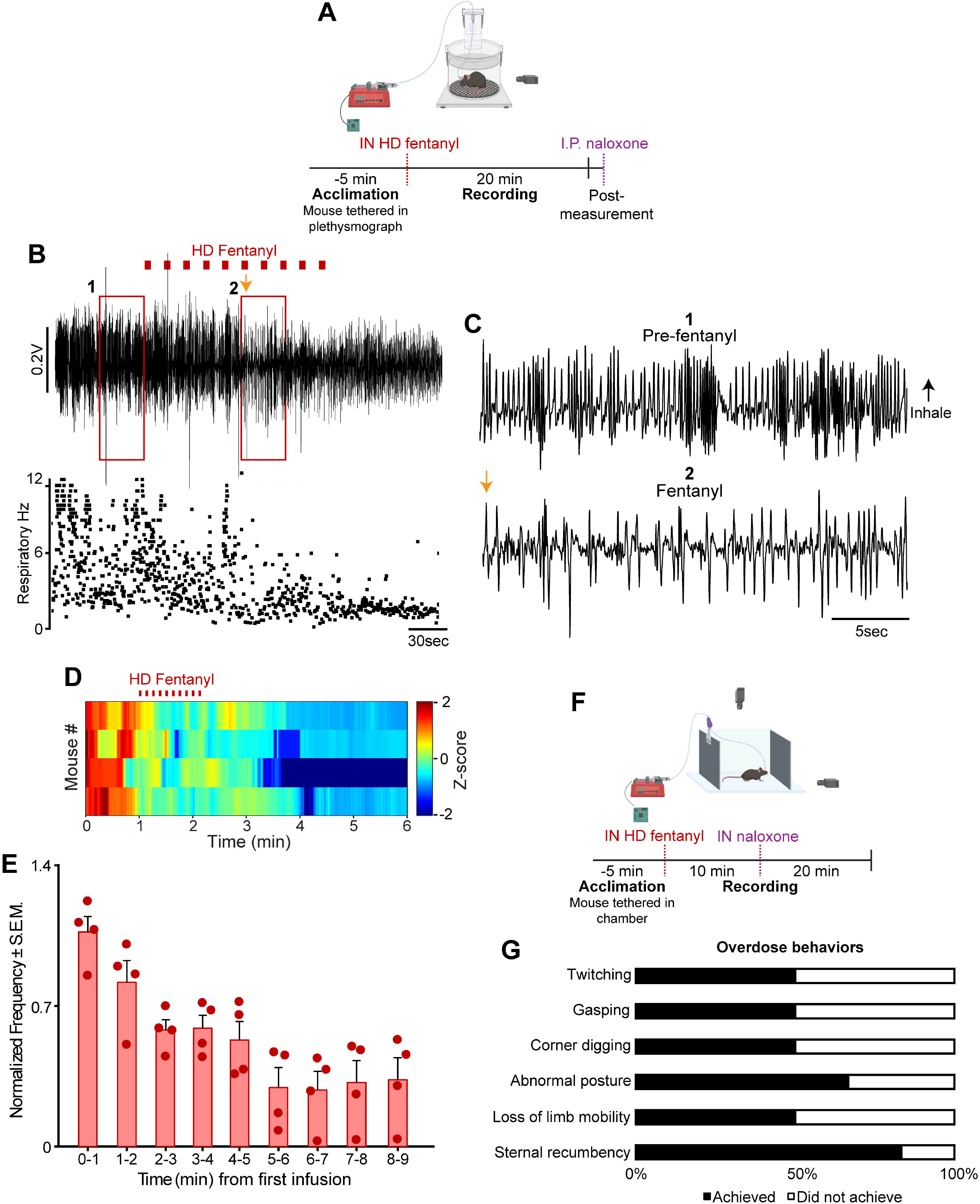
Overdose behavioral and physiological symptoms upon administration of high dose IN fentanyl. **A.** Plethysmograph setup for the measurement of respiratory depression at the onset of IN HD fentanyl and experimental timeline. **B.** Respiratory trace and frequency of breaths observed from a mouse during high dose fentanyl administration in the plethysmograph. **C.** Traces from (**B**), showing respiratory dynamics from a period before fentanyl and a period after fentanyl administration. Orange arrows represent the same time point in the compressed and extracted trace, and is occurrence 72sec post fentanyl delivery. **D.** Instantaneous frequency of respiration of all animals when high dose fentanyl was delivered. **E.** Instantaneous frequency of respiration calculated in 60-second bins for high dose fentanyl administration. **F.** Setup for behavioral measurements after IN HD fentanyl and IN naloxone administration and experimental timeline. **G.** Proportion of mice that achieved each behavioral overdose readout during open-field measurement. One animal, due to being a statistical outlier, is not included in data for **D** & **E**. Created in BioRender. Wesson, D. (2027) https://BioRender.com/7xd28hz.

When IN HD fentanyl was administered to the mice in the open field chamber (**Fig. 5F**), canonical opioid overdose behaviors^45–50^ were observed. 50% of mice demonstrated twitching, gasping, a stereotypy we refer to as “corner digging,” and loss of limb mobility (ataxia). 83.3% achieved sternal recumbency, and 66.7% achieved an abnormal, postural flexion, posture (**Fig. 5G**). These results highlight the respiratory consequences of IN HD fentanyl in mice.

### Rapid recovery of overdose after IN naloxone

To measure respiratory recovery after opioid overdose, the other half of the mice (as described above) received an I.P. injection of HD fentanyl and received IN naloxone in the whole-body plethysmograph in a single testing session (**Fig. 6A**). We found a robust and rapid rescue of respiratory depression, with breathing recovering after naloxone administration (**Fig. 6B**). Alert respiratory dynamics, including sniffing, were seen at about 85 seconds post-first infusion, demonstrating the beginning of respiratory rescue (**Fig 6C**). There was a consistent increase in respiratory frequency after the administration of IN naloxone, with full recovery seen at around 180 seconds after the first infusion (**Fig. 6D**). Progressive increases in breathing frequency were observed across the population of mice following IN naloxone, with respiration being significantly recovered as soon as 3 minutes post-infusion start (Repeated measures ANOVA, *F*(8,24) = 27.843, *p* < 0.0001; 0 min vs 3 min: *p* = 0.038; **Fig 6E**).

**Figure 6.**
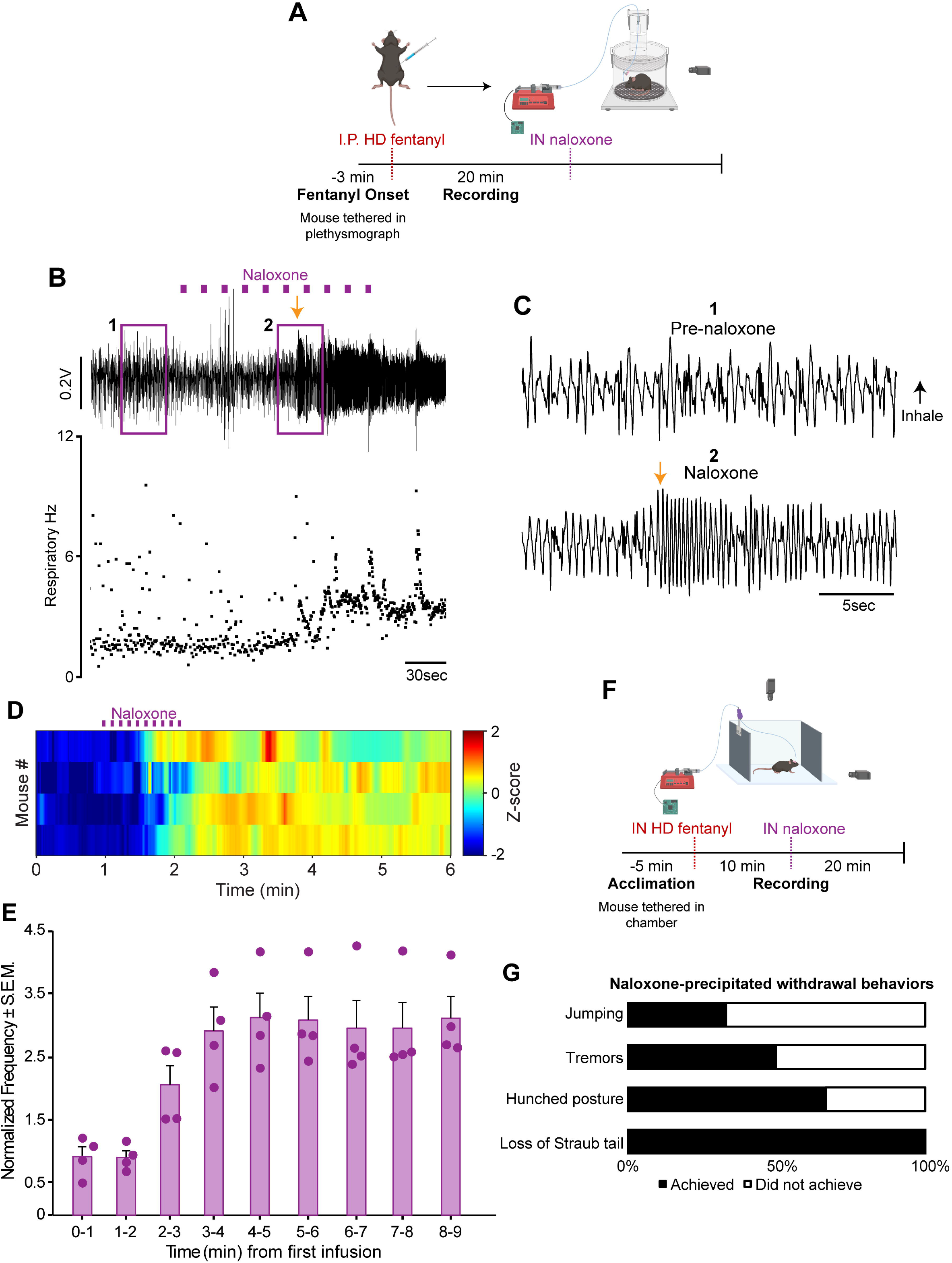
Rapid recovery of overdose upon delivery of IN naloxone. **A.** Plethysmograph setup for the measurement of respiratory recovery at the onset of IN naloxone after fentanyl-induced respiratory depression and experimental timeline. **B.** Respiratory trace and frequency of breaths observed from the same mouse from (**5B**) during naloxone administration in the plethysmograph after being in respiratory depression. **C.** Traces pulled out from (**B**), showing respiration from a period before naloxone and a period after naloxone administration. Orange arrows represent the same time point in the compressed and extracted trace, and occurrence 85sec post IN naloxone delivery. **D.** Instantaneous frequency of respiration of all animals when IN naloxone was delivered after the onset of respiratory depression via I.P. fentanyl. **E.** Instantaneous frequency of respiration calculated in 60-second bins for administration of IN naloxone after the onset of respiratory depression via I.P. fentanyl. **F.** Setup for behavioral measurements after IN HD fentanyl and IN naloxone administration and experimental timeline. **G.** Proportion of mice that achieved each behavioral recovery readout during open-field measurement. One animal, due to being a statistical outlier, is not included in data for **D** & **E**. Created in BioRender. Wesson, D. (2027) https://BioRender.com/7xd28hz.

Mice that received IN HD fentanyl in the behavioral chamber (as described above) then received IN naloxone (**Fig. 6F**). All mice displayed a loss of Straub tail at the onset of IN naloxone. 66.7% of animals achieved hunched posture, 50% achieved tremors, and 33.3% demonstrated jumping behavior (**Fig. 6G**). These results point towards the expected, but not before shown, ability for IN naloxone to recover mice from an overdose of HD fentanyl.

## Discussion

Here, we conducted two separate experiments to examine the impact of IN opioids on DA and opioid dynamics in the brain, and, in separate mice, their impact on behavior, including respiratory depression. While extensive work has examined the behavioral impact of opioids, including morphine and fentanyl following their delivery in the periphery (via S.C., I.P., I.V., or oral delivery), no prior work has examined their impact following IN administration in mice. Likewise, the impact of IN opioids on DA and opioid signaling has not been reported. Our results indicate that IN opioids impact opioid-related behaviors in manners that mimic those following opioid delivery through S.C. or I.P. routes, that IN naloxone can overcome opioid overdose, including opioid induced respiratory depression, and that IN opioids dynamically sculpt neural signaling. Collectively, the results herein help to establish a model for studying the mechanisms underlying opioid use disorder in a manner which recapitulates IN opioid use.

### Physiological outcomes of IN opioids

To determine physiological outcomes of IN opioids, we used fiber photometry to monitor DA and opioid dynamics in the ventral striatum and, in separate experiments, respiratory behavior. By being able to deliver IN opioids while simultaneously recording respiratory changes or monitoring neural dynamics without physically disturbing the mice, we present novel information on the real-time changes of opioid-induced respiratory depression, respiratory recovery, and population-level neuromodulator/neuropeptide activity, specifically following IN opioid delivery.

Opioids act upon opioid receptors in the ventral tegmental area to modulate the activity of postsynaptic DAergic neurons and thereby form a circuit for the modulation of DA release in downstream brain regions, including the NAc^18–22,51^. How these dynamics may occur upon IN opioids, if at all, has remained unknown. As anticipated by the above actions of opioids on disinhibiting ventral tegmental area DAergic neurons^52^, here we uncovered a prompt and prolonged increase in mesolimbic DA, indicated by rises in rDA3.8m emission amplitude, by both IN morphine and fentanyl. We further showed upon IN-achieved opioid receptor antagonism by delivery of naloxone, that there is a restoration of DA to pre-opioid overdose levels (**Fig 1**). We predict based upon prior *ex vivo* work^52^, that is due to the direct actions of naloxone on mu opioid receptors in the ventral tegmental area. The rise in mesolimbic DA levels demonstrates that morphine and fentanyl, when administered IN, are rapid and effective in reaching the canonical reward system. Recent work monitoring noradrenergic neurons in the locus coeruleus uncovered that fentanyl results in brief phasic events in activity^53^. While the route of fentanyl administration was different, that finding together with ours suggests, not surprisingly, that fentanyl differentially impacts neuromodulatory centers throughout the brain.

Simultaneous to the rise in NAc DA, IN morphine and fentanyl also each resulted in a short-lasting increase in NAc opioid dynamics, as indicated by an increase in the deltaLight emission amplitude. We hypothesize that these dynamics are due to naloxone binding the mutant opioid receptor (deltaLight) as a competitive antagonist^54^, resulting in fluorescence emission. This rise in the deltaLight signal is, however, short-lived (**Fig 1**), which we predict is due to the eventual displacement of naloxone. That NAc DA levels indicated by rDA3.8m, but not deltaLight levels, returned to or near pre-opioid overdose levels following naloxone, and around the time wherein behavior indicated overdose recovery, may point towards a mechanism of importance for opioid overdose. Specifically, we predict that these findings implicate a likely important role for mesolimbic DA, specifically DA levels and actions upon postsynaptic DA receptor expressing neurons in the ventral striatum, in mediating opioid overdose reversal since DA was the only of the two measures which returned to pre-overdose levels.

As an additional physiological measure, we observed that when the mice received IN morphine and LD fentanyl, they exhibited mild respiratory depression. Breathing frequency during IN LD fentanyl and morphine was significantly different than that of saline control as soon as 2 minutes post-infusion (**Fig 4**). When IN HD fentanyl was delivered, the mice displayed greater respiratory depression, with breathing frequency decreasing steadily and rapidly during and after administration (**Fig 5**). As expected, IN naloxone rescued opioid induced respiratory depression (**Fig 6**). While these outcomes are not unlike prior reports following I.P opioid delivery, here by IN delivery, we found the short latencies by which IN opioids exert their effects, which points towards the rapid distribution of opioids into brain regions responsible for opioid induced respiratory depression by IN administration, including those in the respiratory brainstem^32,35^.

### Behavioral outcomes of IN opioids

In a separate series of experiments, we monitored several behaviors well-known to be impacted upon opioid delivery in rodents. We were most interested in answering two specific questions: First, do the same types of behaviors occur upon IN opioids as reported by prior work following S.C. or I.P. opioids? Second, given the accessibility to the brain from the IN route, do these behaviors occur more rapidly following IN opioid delivery? Following I.P. or S.C. opioid delivery, mice and rats display stereotyped behavioral responses. These include the Straub tail response^30,31,55^, darting, hyperlocomotion, and tip-toeing^37–39^ upon morphine and LD fentanyl. These also include overdose behaviors upon HD fentanyl administration, including sternal recumbency^45^, gasping^47^, twitching^46^, loss of limb mobility (ataxia)^49^, “corner digging” stereotypy^48^, and abnormal posture (postural flexion)^50^. Opioid receptor antagonism with naloxone also elicits specific behaviors including tremors, hunched posture, loss of Straub tail, and leaping^56–58^. Through IN delivery of opioids, we observed all of the behaviors noted above from studies delivering opioids I.P. or S.C. (**Fig 3**). This answers our first primary question, and indicates that the route of administration is not a determinant factor in the recruitment of CNS and likely also the PNS circuits needed for the orchestration of those opioid-induced behaviors. Moreover, these results support the use of IN opioid delivery to study behavioral effects of opioids. Through IN opioid delivery, we were also able to assess the real-time changes in behavior not previously possible following S.C. or I.P. delivery, and thereby answer our second question regarding the latency of opioid-induced behavioral changes. This is because we could deliver opioids without manually intervening with the mice, which would fundamentally interrupt ongoing behaviors, induce a stress response, etc. Through these real-time uninterrupted observations, we found that with IN morphine, latency to Straub tail response occurred in less than three minutes post-infusion and that in some extreme cases, after IN LD fentanyl, the Straub tail response would occur within just one minute following infusion (**Fig 3**). Previous reports indicated that S.C. injections of morphine elicit a Straub tail response within 3-6 minutes^59^ and I.P. injections of fentanyl elicit a response within 1-5 minutes^60^. Thus, our results suggest that IN opioids may activate neural systems more rapidly than following S.C. or I.P. delivery in manners resulting in prompt behavioral changes. We predict this outcome holds implications for other effects of IN opioids, including possibly the psychoactive effects of IN opioids. For instance, if these behavioral effects may arise in part by influences of opioids on glutamatergic signaling and/or DAergic signaling, our results may support that IN opioids can more rapidly trigger changes in glutamate or DA – two systems integral for the reinforcing effects of opioids^3,61^.

Interestingly, while not a main goal of our study, we observed a significant decrease in facial grooming, rearing, and itching upon IN morphine and LD fentanyl (**Fig 3**). Prior research point towards reductions in these specific behaviors as being due to activation of kappa opioid receptors^62^ and also that activation of DA D3 receptors in the ventral striatum coordinate the occurrence of facial grooming^63^. This unexpected finding that IN opioids decrease grooming therefore suggests the possible access and recruitment of ventral striatum D3 receptor expressing neurons upon IN opioid delivery – an outcome which could hold translational importance given the role of D3 receptors in a variety of psychiatric conditions^64^.

### Conclusions

Taken together, through these studies examining changes in DA and opioid dynamics in the NAc, as well as behavior upon IN opioid delivery, we have uncovered some similarities, but also differences, in the outcomes of IN opioids to those reported in the literature following other routes of administration. More importantly, our results highlight the rapid action of IN opioids upon opioid and mesolimbic DA dynamics, as well as on specific behaviors. Moreover, our results point towards a likely important role for mesolimbic DA for overcoming opioid overdose. Now knowing the fundamental actions of IN opioids on several aspects of physiology and behavior, this work may serve as a foundation for future studies using this IN opioid delivery approach, combined with measures of opioid reinforcement in mice, to uncover novel mechanisms underlying the reinforcing effects of IN opioids. Based upon the many differences availed by the IN route of administration, we hypothesize those insights will both diverge from, and more importantly, better bolster, our in-depth understanding for the mechanisms mediating opioid reinforcement.

## Data availability

The authors will provide original full data upon reasonable requests.

## Code availability

No custom code was used in this study.

## Acknowledgements

This work was supported by R01DA049449 and R21DA063991. K.J.E. is supported by T32DC015994. A.G.V. is supported by R00HL159232. M.F.R received support from T32DC015994 and F31DA066224. We thank Ellyse Thomas for her expert assistance with animal care. We also thank Dr. Zihao Zhang for his expertise in photometry data analysis.

## Author Contributions

Conceptualization: KJE, DWW

Methodology: KJE, MFR, AGV, DWW

Investigation: KJE, DWW

Visualization: KJE, DWW

Funding acquisition: KJE, DWW, AGV, MFR

Supervision: DWW

Writing – original draft: KJE, DWW

Writing – review & editing: KJE, MFR, AGV, DWW

